# The architecture of membrane structures involved in hepatitis C virus genome replication revealed in close-to-native conditions by cryo-electron tomography

**DOI:** 10.1101/2025.04.24.650446

**Authors:** Upasana M. Sykora, Thomas J. O’ Sullivan, Yehuda Halfon, Juan Fontana, Mark Harris

## Abstract

Hepatitis C virus (HCV) infection induces extensive rearrangements of host cytoplasmic membranes, leading to the formation of multiple membranous structures that facilitate RNA replication. Current knowledge of these membranous structures has largely relied on correlative light and electron microscopy (CLEM) techniques using chemical fixation and resin embedding. To overcome these limitations, cryo-preserved cells were prepared using cryo-focused ion beam (cryo-FIB) milling and cryo-ultramicrotomy. For the first time, the contents within the membranous structures have been observed *in-situ* using cryo-electron tomography (cryo-ET) performed on lamellae (prepared via cryo-FIB) and on ultrathin sections (prepared via cryo-ultramicrotomy) from HCV subgenomic replicon harbouring cells. Observations from 112 cryo-electron tomograms of cryo-FIB-derived samples revealed the presence of densities within the inner vesicles of a subset of single-, double-membrane vesicles (SMVs, and DMVs respectively), as well as within multi-vesicular bodies (MVBs), which might represent the viral replication machinery. Notably, this study represents the first direct visualisation of the arrangement of non-structural proteins within a multi-membrane vesicle (MMV) observed from cryo-electron microscopy of vitreous sections (CEMOVIS). The cryo-ET methodologies established here lay the groundwork for future investigations into the architecture of the HCV replication complex, leveraging advanced computational tools for deeper structural and functional analysis.

## Introduction

Hepatitis C virus (HCV) is a positive sense RNA virus that primarily infects hepatocytes in the liver, leading to both acute and chronic hepatitis. The latter results in progressive liver damage, including cirrhosis, liver failure, and hepatocellular carcinoma (Martinello et al., 2023). In common with other positive-sense RNA viruses, HCV induces cytoplasmic membrane rearrangements in the host cell to form specialised structures that support genome replication (Wolff et al., 2020). Current ultrastructural insights into host membrane rearrangements during HCV infection have primarily come from fluorescence microscopy (FM), electron microscopy (EM), and soft X-ray microscopy (Ferraris et al., 2010; Romero-Brey et al., 2012; Paul et al., 2014; Lee et al., 2019). For example, these studies have shown that the membrane rearrangements arise from the ER and autophagy pathways (Mohl et al., 2016; Pérez-Berná et al., 2016). These membranes form vesicular networks, observed during both viral infection and after transfection with subgenomic replicons (SGR), which are often referred to as viral replication factories or the membranous web (MW) (Gosert et al., 2003; Moradpour et al., 2003; Ferraris et al., 2010). The MW comprises various virus-induced vesicles, and includes single, double, and multi-membrane vesicles (SMV, DMV, MMV, respectively), multivesicular bodies (MVBs) and lipid droplets (LDs) (Romero-Brey et al., 2012). These structures have been proposed as potential replication organelles (ROs), serving as platforms for the replicase machinery.

HCV RNA replication is orchestrated by the five non-structural proteins NS3, NS4A, NS4B, NS5A and NS5B, which are necessary and sufficient to form MW in the cytoplasm of infected or transfected cells (Lohmann et al., 1999; Gosert et al., 2003). NS3/4A functions as a protease and helicase, facilitating HCV polyprotein cleavage and the unwinding of RNA secondary structures (Bartenschlager et al., 1995). NS4B contributes to MW formation alongside NS5A (Moradpour et al., 2003). NS5A is a multifunctional viral protein involved in viral replication and assembly (Hughes et al., 2009); notably it binds NS5B and modulates its RNA-dependent RNA polymerase activity (Shirota et al., 2002). Specifically, within the MW, DMVs have been primarily hypothesised as the main HCV replication site through immunogold labelling of NS3 and NS5A. Thus far antibodies against NS5B have not been used to designate a specific vesicle by immunogold labelling. Additionally, double-stranded RNA (dsRNA) (a marker for RNA replication) showed colocalisation with NS5A by immunofluorescence and correlated with the production of DMVs over the course of infection, further supporting DMVs as active sites of HCV RNA synthesis (Romero-Brey et al., 2012). Furthermore, another study employed an SGR containing an NS5A-mCherry fusion protein and suggested that the majority of NS5A-positive regions (∼35%) were in DMVs and MMVs (Grünvogel et al., 2018). However, the same studies also showed that immunogold labelling of NS3 and NS5A also localised to SMVs, LDs and ER (Romero-Brey et al., 2012), and that a relevant proportion of NS5A-mCherry regions corresponded to MVBs (∼20%), ER (∼7%), LDs (∼7%) and mitochondria (∼5%) (Grünvogel et al., 2018). Additionally, chronically infected liver tissue samples from patients does not present any DMVs, and the MW appeared primarily as clusters of SMVs closely associated with ER and LDs (Blanchard and Roingeard, 2018). Overall, although DMVs are suggested as the primary ROs for HCV, the type of membranous structure utilised as the major HCV replication site is still inconclusive.

To date, the EM methods employed to study cellular architecture following infection (or in cells harbouring an SGR) have yielded insights into membrane rearrangements and the roles of individual NS proteins in these processes (Ferraris et al., 2010; Romero-Brey et al., 2012). However, the sample preparation methods for visualisation by EM utilised chemical fixation, high pressure freezing with freeze substitution, resin embedding and sectioning techniques. Therefore, we explored the recent advancements in cryogenic sample preparation techniques and cryo-electron tomography (cryo-ET), which facilitate imaging in close-to-native conditions, to gain insight into the ultrastructure of these membranous structures in cells harbouring SGR. Overall, our results confirm the presence of all the above-mentioned vesicles within the HCV MW, and supports a model in which internal densities, potentially corresponding to viral and/or cellular components, accumulate inside inner vesicles (InVs), present inside both DMVs and SMVs, and in which NS5A-eGFP accumulates around MMVs within cells stably harbouring an HCV SGR.

## Materials and Methods

### Genome sequences

The HCV genome sequence for JFH-1 (genotype 2a) used in the analysis was sourced from GenBank with accession number AB047639.1. The JFH-1 SGR (Wakita et al., 2005) was modified to replace the firefly luciferase (FF-luc) reporter with a fusion of FF-luc and neomycin phosphotransferase (termed Feo) to create the pSGR JFH-1 Feo NS3-5B construct (Wyles et al., 2009; Ross-Thriepland et al., 2015) and a green fluorescent protein (GFP) was inserted in NS5A-domain III (using insertion site P418) (Moradpour et al., 2004). The final construct used for this study was SGR JFH-1 Feo NS3-5B (NS5A-eGFP), hereafter referred to as SGR-NS5A-eGFP.

### Linearisation of plasmid DNA for generating *in-vitro* RNA transcripts

Plasmid DNA (∼5 µg) was digested with XbaI (NEB) in CutSmart buffer (NEB) at 37 °C for 1 hour and purified by phenol-chloroform extraction. The purified linearised DNA (∼1 µg) was used for RNA synthesis using Ribomax Express T7 kit (Promega) and purified using an RNA clean and concentrator kit (Zymo Research). The RNA transcript integrity was analysed on a 1% agarose gel, and concentration was measured using a Nanodrop spectrophotometer.

### Cell culture

Huh7 cells were maintained in Dulbecco’s modified Eagle’s medium (DMEM, Sigma) supplemented with 10% foetal bovine serum (FBS, Gibco), 50 Units/ml penicillin, 50 μg/ml streptomycin, 1% nonessential amino acids (NEAA, Lonza) and 2.8% HEPES (Lonza). Cells were maintained in a humified incubator at 37 °C with 5% CO_2_. To generate stable SGR-NS5A-eGFP harbouring Huh7 cells, 2 µg of replicon RNA was electroporated into 2 × 10^6^ cells using a square-wave protocol (260 V, 25 ms). Electroporated cells were mixed with non-electroporated cells to a final density of 1.8 × 10⁵ cells and seeded in 6-well plates. After 48 h of incubation, cells were washed with PBS and subjected to selection with 700 µg/ml G418. Following multiple selection rounds, the G418 concentration was reduced to 400 µg/ml for maintenance of the polyclonal cell population.

### Fluorescence microscopy

To prepare coverslips for confocal fluorescence microscopy, 22 mm coverslips were cleaned with dH₂O, treated with 1M HCl, washed, incubated in ethanol, dried, and autoclaved. Mock or SGR-expressing cells (5 × 10⁴) were seeded on coverslips or sorted by FACS for fluorescence-based selection. Sorted cells were incubated for ∼21 h, fixed with 4% formaldehyde, permeabilized with 0.2% Triton X-100, and blocked with 3% BSA-PBS. Coverslips were then stained with BODIPY dye (1:1000), washed, and mounted with ProLong Gold Antifade reagent with DAPI. Mounted slides were incubated for 2.5 h at room temperature and stored at 4 °C.

Z-stack confocal microscopy images were captured using a Zeiss LSM880 upright confocal microscope. After acquisition, the images were processed with Fiji software (Schindelin et al., 2012). Wide-field fluorescent images were acquired with the EVOS microscope (Thermo Fisher Scientific).

### Resin embedding and sectioning for cellular EM

Huh7 cells (control and stably expressing SGR-NS5A-eGFP) were trypsinised and pelleted. The resulting cell pellet was fixed in with 2.5% glutaraldehyde (EM Grade, Agar Scientific) in 0.1M sodium phosphate buffer for at least 2.5 h at room temperature. The cell pellet was washed twice for 30 min in 0.1M sodium phosphate buffer, post-fixed with 1% osmium tetroxide in 0.1M phosphate buffer for 1 h and washed again twice for 30 min with the same buffer. Dehydration was achieved by incubating the pellet for 20 min in increasing concentrations of ethanol: 20%, 40%, 60%, 80%, and twice in 100%. The pellet was then treated with propylene oxide twice for 20 min each. A freshly prepared resin mixture of 50% propylene oxide and 50% Araldite epoxy resin was then added to the pellet and allowed to infiltrate for several h to overnight (Luft, 1961). This was followed by infiltration with a 25% propylene oxide and 75% Araldite epoxy resin mixture for 3 h, and with 100% Araldite epoxy resin for 3-8 h. Polymerization was then carried overnight at 60 °C. Ultrathin sections (80-100 nm) were collected on slotted copper EM grids with formvar support and stained with saturated uranyl acetate for 2 h and Reynolds lead citrate for 15 min.

### Image acquisition using the FEI Tecnai T12 TEM

Cellular EM images from ultrathin sections prepared through resin embedding and sectioning were captured using the FEI Tecnai T12 TEM operated at 120 kV.

### Cell culture workflow on EM grids

For cryo-ET, Quantifoil Finder 200 mesh gold grids R1.2/1.3 (Gilder G200F1) and C-flat 200 mesh gold 1.2/1.3 (Electron Microscopy Sciences) were utilised. The grids were glow discharged in a Quorum GloQube (Quorum Technologies) to enhance hydrophilicity and impart a negative charge. A negative polarity cycle of 20 mA was applied for 30-60 seconds.

Next, the grids were positioned in the centre of a glass-bottom cell culture dish (Nunc TM, 150680). Grids were functionalised with 7 μl of fibronectin (Sigma-Aldrich) at a concentration of 25 μg/ml and incubated at 37 °C for 1 h. Concurrently, fluorescence-activated cell sorting (FACS) was performed to collect 5 × 10^3^ and 10 × 10^3^ cells and were pelleted and resuspended in 50 μl of complete DMEM before being seeded onto the EM grids at various cell densities. Following a 20 min incubation at room temperature, 2 ml of complete DMEM was added to each dish, and the cells were incubated at 37 °C with 5% CO_2_ for 20-24 h to promote optimal cell attachment and spread.

### Plunge freezing of cells on EM grids

After culture, the cells on the EM grids were plunge-frozen using a Leica EM GP Automatic plunge freezer (Leica Microsystems) in liquid ethane at -180 °C and transferred in liquid nitrogen. To facilitate blotting, 3 μl of 1X PBS was applied to the side of the grid containing the cells, and 0.5 μl was applied to the back of the grid within the humidifier chamber. The chamber was maintained at 90% humidity and 8 °C, with blotting times adjusted between 5 and 8 seconds for each sample. The grids were blotted using No.1 Whatman paper, then stored in liquid nitrogen. Subsequently, clipped into C-rings and placed into autogrids for screening and data collection compatible with the FEI Titan Krios TEM (Thermo Fisher Scientific).

### Sample thinning by cryo-focussed ion milling-scanning EM (cryo-FIB-SEM)

The lamellae generation was carried out at the Electron BioImaging Centre (eBIC) at Diamond Light Source, utilising the Scios cryo-FIB-SEM (Thermo Fisher Scientific) and Aquilos cryo-FIB-SEM (Thermo Fisher Scientific). Lamellae were prepared using a focused gallium ion beam on a dual-beam focused ion beam-scanning electron microscope (FIB-SEM) at a stage temperature of -180 °C. Cells located in the centre of the grid squares were selected for milling. To achieve uniform milling of the cells during the process, an organo-metallic platinum layer was applied prior to milling. This was followed by a thin layer, or splutter coat, of Pt to enhance SEM imaging. The rough milling was conducted in three gradual steps at a stage angle of 12° using a gallium beam, followed by a final fine milling step to achieve the desired nominal thickness of the lamellae. The final beam current used when milling was performed on SCIOS was 37 pA, while on Aquilos was 50 pA. Progress was monitored using SEM on the SCIOS system (operating at 2.2-10 kV with ETD and T2 detectors) and on the Aquilos system (operating at 2-5 kV with ETD detectors and Auto TEM).

### Sample vitrification by high-pressure freezing

SGR-NS5A-eGFP cells were vitrified using a high-pressure freezing method. Approximately 1 × 10^6^ cells per ml were resuspended in 100 µl of cryoprotectant (20% w/v Dextran 40,000 in DMEM). The sample was placed into the 100 µm deep wells of an Au type-A specimen carrier (Leica, catalogue no.16770152), ensuring the wells were slightly overfilled to prevent air bubbles, and then covered with the flat side of a lipid-coated (L-⍺-Phosphatidylinecholine, Sigma, 61771) Au type-B carrier (Leica, catalogue no.16770153). Excess liquid was absorbed with a filter paper, and the sample was then high-pressure frozen at approximately 2100 bar and -190 °C for 300 ms using a Leica EM ICE High-Pressure Freezer (Leica Microsystems).

### Cryogenic fluorescence microscopy

High-pressure frozen, cryo-preserved carriers and cryo-EM grids containing SGR-NS5A-eGFP cells were screened and imaged using a Leica THUNDER Imager EM cryo CLEM (Leica Microsystems) equipped with a HC PL APO 50x/0.9 NA cryo objective, an Orca Flash 4.0 V2 sCMOS camera (Hamamatsu Photonics), and a Solar Light Engine (Lumencor). Z-stack images were acquired with a frame size of 2048 x 2048 pixels, at 30% intensity and an exposure time of 0.2 seconds, utilizing the LASX software (Leica Microsystems). The resulting images were processed using Fiji software (Schindelin et al., 2012).

### Sample thinning by cryo-ultramicrotomy

High pressure frozen samples were sectioned at cryogenic temperatures (-160 °C) using a Leica UC7 equipped with an FC7 chamber, trimming (Diatome, Trim20), and cryo-sectioning (Diatome, cryo immune, 3 mm) Diamond knifes, and micro-manipulators (Studer et al., 2014). Images were captured with a stereomicroscope inside the FC7 chamber and correlated with fluorescent images obtained from the Leica Thunder Imager (Leica Microsystems) to precisely identify the regions for cryo-ET. A trapezoidal block of tissue measuring 150 × 100 x 40 µm was shaped around fluorescent targets, from which ribbons of either 70 or 40 nm thickness were produced. These ribbons were then adhered to Quantifoil R2/2, Cu 300 mesh grids (EMS) using an electrostatic gun, after the grids had been glow discharged for 60 seconds at 30 mA by glow discharge (Easy Glow, Cressington).

### Image correlation of cryo-EM and cryo-FM maps

Maps software (Thermo Fisher Scientific) was utilised to correlate fluorescent images on-the-fly at the TEM. Low magnification electron micrographs (atlas map: 125X, pixel size 819.2 Å, spot size: 7, illuminated area: 1100 µm) were aligned based on broken grid squares and distinctive patterns embedded in the centre of Quantifoil grids. At medium magnification (overview: 580X, pixel size 224.4 Å, spot size: 6, illuminated area: 313 µm), specific grid squares were matched. At high magnifications (search map: 8700X, pixel size 28.8 Å, spot size: 10, illuminated area: 12 µm), holes in the carbon support film served as fiducial markers to facilitate correlation. The correlation between the fluorescent images and electron micrographs after tomogram collection was achieved using the Matlab script Correlate.tex (Schorb and Briggs, 2014).

### Cryo-ET

Cryo-ET was conducted using the FEI Titan Krios TEM (Thermo Fisher Scientific), operated at 300 kV and equipped with a Falcon 4i direct electron detector and a Selectris imaging filter (Thermo Fisher Scientific) in energy-filtered TEM (EFTEM) nanoprobe mode. Two different acquisition schemes were employed for imaging lamellae. The first one, involved a magnification of 42,000 X, corresponding to a pixel size of 3.0 Å at the specimen level, using Tomography 5.11 or 5.13 software (Thermo Fisher Scientific). A dose-symmetric acquisition scheme (Hagen et al., 2017) was implemented from - 60° to 60° in 2° increments, maintaining a constant electron dose of ∼1.5 e−/Å² per projection, corresponding to a total dose of ∼95 e−/Å². The target defocus was between -3 and -5 μm and an energy filter slit set to 10 eV was used. Individual projections were captured in counting mode through dose fractionation. Four frames per projection were aligned and summed ‘on-the-fly’ using custom Python scripts that utilized commands from the MotionCor2 (Zheng et al., 2017) and IMOD (Kremer et al., 1996) software packages. The second acquisition scheme was similar to the first one, with the following differences: the magnification employed was 64,000 X, with a corresponding pixel size of 1.9 Å. The electron dose was ∼2.9 e−/Å² per projection, corresponding to a total dose of ∼120 e−/Å². The tilt range covered was from 71.2° to -46.8° in 3° increments, starting at an angle of 12.14°. Eight frames per projection were collected.

Similarly, tilt series for the cryo-ultrathin sections were collected at a magnification of 53,000 X, with a corresponding pixel size of 2.40 Å. A dose-symmetric acquisition method was employed from -60° to 60° in 3° increments, maintaining a constant electron dose of ∼2.2 e−/Å² per projection, resulting in a total electron dose of ∼91 e−/Å². The target defocus ranged from -3 to -5 μm, and an energy filter slit of 10 eV was utilized. Six frames per projection were collected.

### Tomogram reconstruction

Raw images were motion corrected using MotionCor2 (Zheng et al., 2017). The tilt series images were aligned and reconstructed via the eTOMO interface of the IMOD software package (Kremer et al., 1996). For lamellae dataset 1, tomograms were generated using fiducial-less alignment. All tomograms were reconstructed with weighted back-projection. For lamellae dataset 2 and the tomograms of cryo-ultrathin sections, reconstruction was carried out using Aretomo software (Zheng et al., 2022). All tomograms were binned 4 times. Tomograms from datasets 1 and 2 are presented after filtering with 3D Gaussian blur function in Fiji. For the CEMOVIS dataset, images are presented without filtering.

### Statistical analysis on diameter of the vesicles

To evaluate the size of each membranous structure, identifiable vesicles and cellular features were summarized for each tomogram based on the existing literature. The data on the lamellae were collected without specific correlative guidance for the NS5A-eGFP foci, leading to the analysis of all measurable membrane structures (SMV, DMV, MMV, MVB, LD) across 112 tomograms (combined dataset). The predominant vesicles visible in the tomograms and measurable included SMVs, followed by DMVs, LDs, MVBs, MMVs, and mitochondria. For each structure, both the minor and major axes were measured using IMOD’s ‘measure’ function with the diameter calculated by averaging these two measurements. The data was exported to GraphPad Prism for presentation.

## Results

### Establishing a system to study HCV MW

Most ultrastructural studies examining HCV MW have utilised the JFH-1 virus or JFH-1-SGR-NS5A-eGFP, supporting its use for structural studies (Ferraris et al., 2010; Romero-Brey et al., 2012; Ferraris et al., 2013; Lee et al., 2019). NS5A plays a critical role in both replication and assembly of the virus and is distributed as puncta throughout the cytoplasm (Eyre et al., 2014), which makes it a key protein to track and study HCV replication.

To validate the use of SGR-NS5A-eGFP (Figure 1A) harbouring cells for subsequent cryo-EM based analysis, the localisation of NS5A-eGFP in proximity to the LDs, as previously reported (Lee et al., 2019), was confirmed by confocal microscopy (Figure 1B). To confirm architectural differences between control Huh7 cells and those stably harbouring the SGR-NS5A-eGFP, resin-embedded cell samples were sectioned, stained, and imaged by transmission electron microscope (TEM) prior to implementing a cryo-electron tomography (cryo-ET) workflow. Overall, noticeable differences were observed in the cellular architecture between control cells and cells harbouring SGR-NS5A-eGFP. Whereas control cells lacked any evidence of intracellular membrane reorganisation, stably harbouring cells contained apparent MWs (Figures 1C and 1D). Therefore, cells stably harbouring SGR-NS5A-eGFP were employed to establish *in-situ* cryo-ET workflows and to gain knowledge into the organisation of the different types of membranous vesicles present within the MW of HCV.

**Figure 1:**
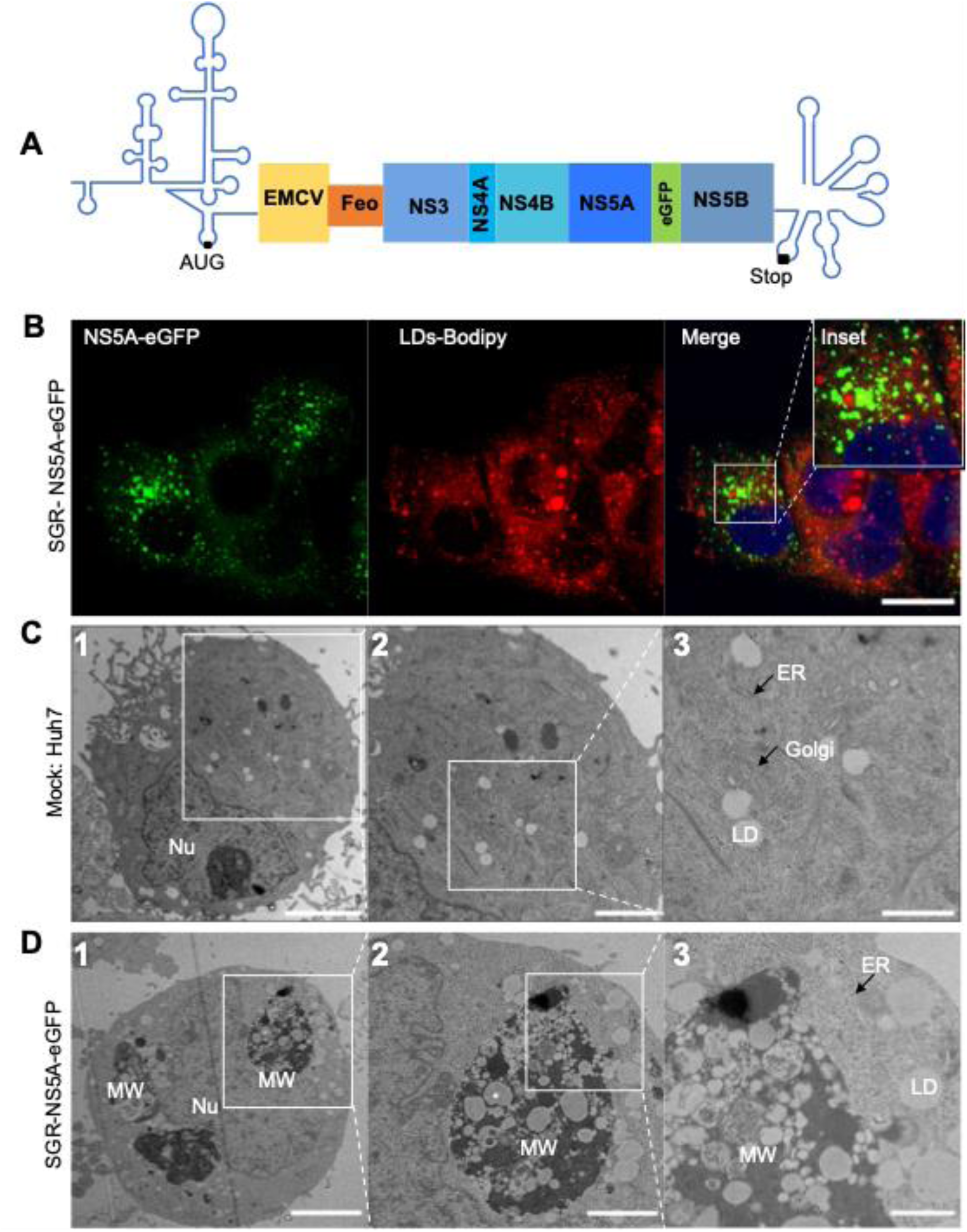
Huh7 stably harbouring SGR-NS5A-eGFP show membrane rearrangements. A) Schematic of the HCV SGR-NS5A-eGFP employed in this study. B) Confocal image of Huh7 cells harbouring SGR-NS5A-eGFP. Cells were fixed with 4% formaldehyde. LDs stained with 1:1000 dilution of BODIPY (558/568) for 1 hour at room temperature and nuclei stained with ProLong gold antifade mountant with DAPI. NS5A-eGFP (green), LDs (red) and nuclei of cells stained with DAPI (blue) are shown. Scale bar: 20 µm. C and D) Representative TEM of ultrathin sections of Huh7 cells (C) and Huh7 cells harbouring HCV SGR-NS5A-eGFP (D). MW: Membranous web, LD: Lipid droplets, Nu: Nucleus, ER: Endoplasmic reticulum. Scale bars: C1,5 µm; D1,4 µm; C2 and D2, 2 µm; C3 and D3, 1µm.

### Sample preparation workflow to generate lamellae by cryo-FIB for cryo-ET

Next, to gain insight into the architecture of the HCV MW in the SGR-NS5A-eGFP harbouring cells in close-to-native conditions, two different approaches were explored: cryo-focused ion beam milling (cryo-FIB) and cryo-electron microscopy of vitreous sections (CEMOVIS).

To establish a cryo-FIB workflow, a homogenous population of cells harbouring SGR-NS5A-eGFP were fluorescently activated cell sorted (FACS) and selected for the brightest eGFP intensities (Figure 2A). These cells were seeded on negatively glow-discharged and fibronectin-functionalised gold grids (Figure 2B). Since cell adherence was variable on each grid, even after functionalisation, optimal cell distribution was analysed by widefield fluorescence microscopy before selecting grids for plunge freezing (Figure 2B and C). Subsequently, grids were screened by cryo-EM for ice thickness and target cells suitable for cryo-FIB milling were selected (Figure 2D). Generally, the success rate of grids with optimum cell targets and ice thickness was ∼ 50% per session. Cryo-FIB was then performed and the presence of LDs in the lamella, as observed through SEM imaging, confirmed that the lamella’s thickness was suitable for cryo-ET imaging (Figures 2E and F).

**Figure 2:**
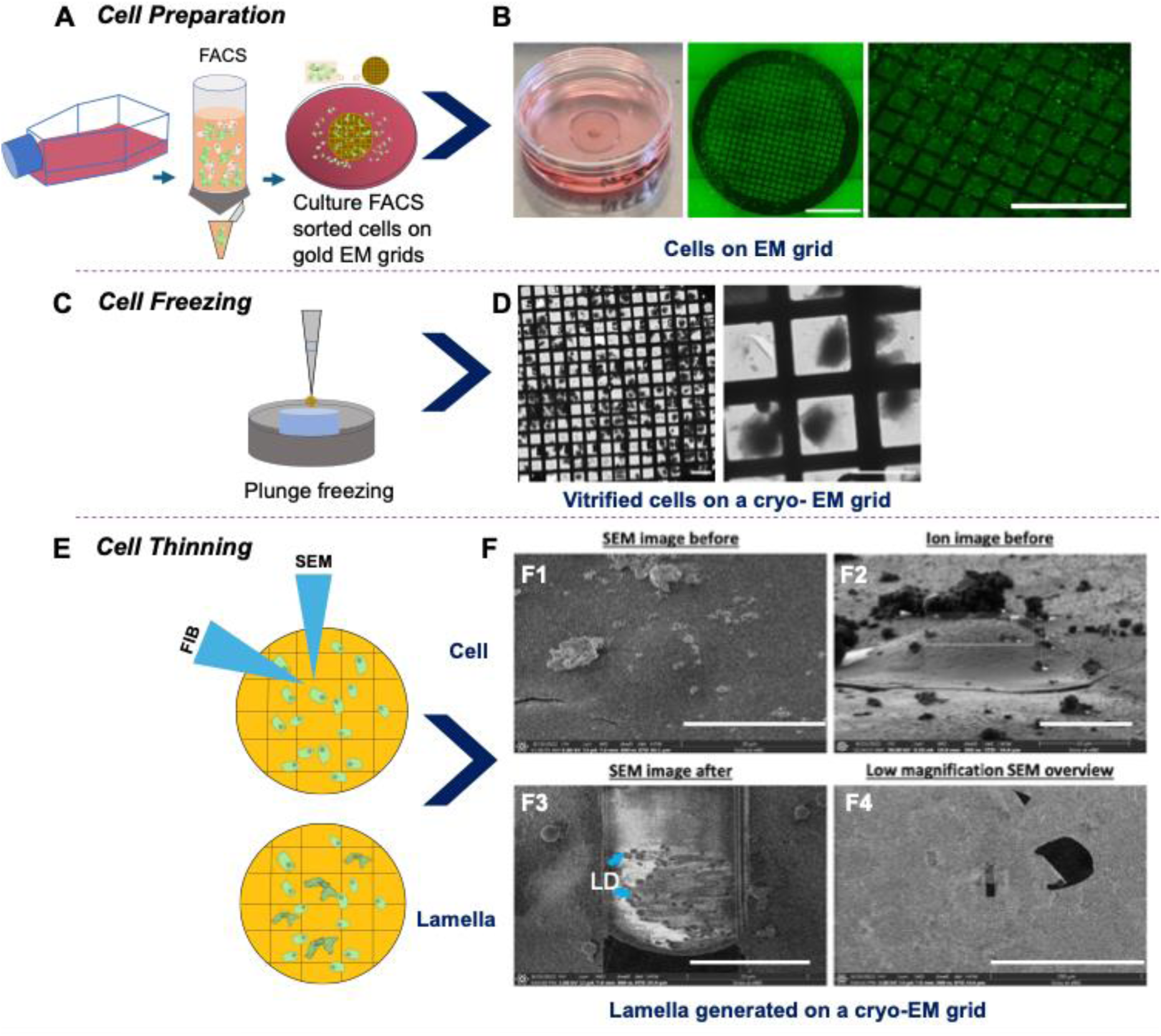
Schematic sample preparation workflow to generate lamellae by cryo-FIB for cryo-ET of Huh7 cells stably harbouring SGR-NS5A-eGFP. Schematics (A, C and E) and images of workflow (B, D and F) to prepare samples for cryo-FIB and cryo-ET imaging of HCV MW. A) Schematic of sample preparation for grid preparation. Cell cultures of Huh7 cells stably harbouring SGR-NS5A-eGFP (left) were FACS sorted (centre) and cultured on glow-discharged and fibronectin-functionalised gold EM grids (right). B) The cell adherence and distribution on the grid was analysed by fluorescence microscopy. Scale bars: 1 mm (centre) and 0.5 mm (right). C) Schematic of plunge freezing process. D) Ice thickness of plunge-frozen grids and identification of cell targets suitable for cryo-FIB were assessed by cryo-EM. Scale bars: 100 µm. E) Schematic of cryo-FIB milling to generate lamellae for cryo-ET. F) SEM images showing the process of cryo-FIB: top, SEM (F1) and ion (F2) images of a target cell prior to cryo-FIB; bottom, SEM image of a lamella in which LDs are apparent (F3) and low magnification SEM image of a lamella (F4). Scale bars: C1, 30 µm; C2 and C3, 10 µm; and C4, 200 µm.

### Quantitative and qualitative analysis of HCV MW using the cryo-FIB/cryo-ET workflow

To investigate the native architecture of HCV the different membranous vesicles within the MW, cryo-ET was applied to the lamellae generated from SGR-NS5A-eGFP harbouring cells. To locate regions for tilt-series acquisition, search maps were utilised (Figure 3A), looking for areas containing MWs. A total of 112 reconstructed tomograms from two datasets were analysed. Among these, 50% contained MVBs, 39% included SMVs, 29% featured DMVs, 15% contained LDs, 12% included ER, 11% featured MMVs, 9% exhibited extensive MW, and 5% contained mitochondria.

**Figure 3:**
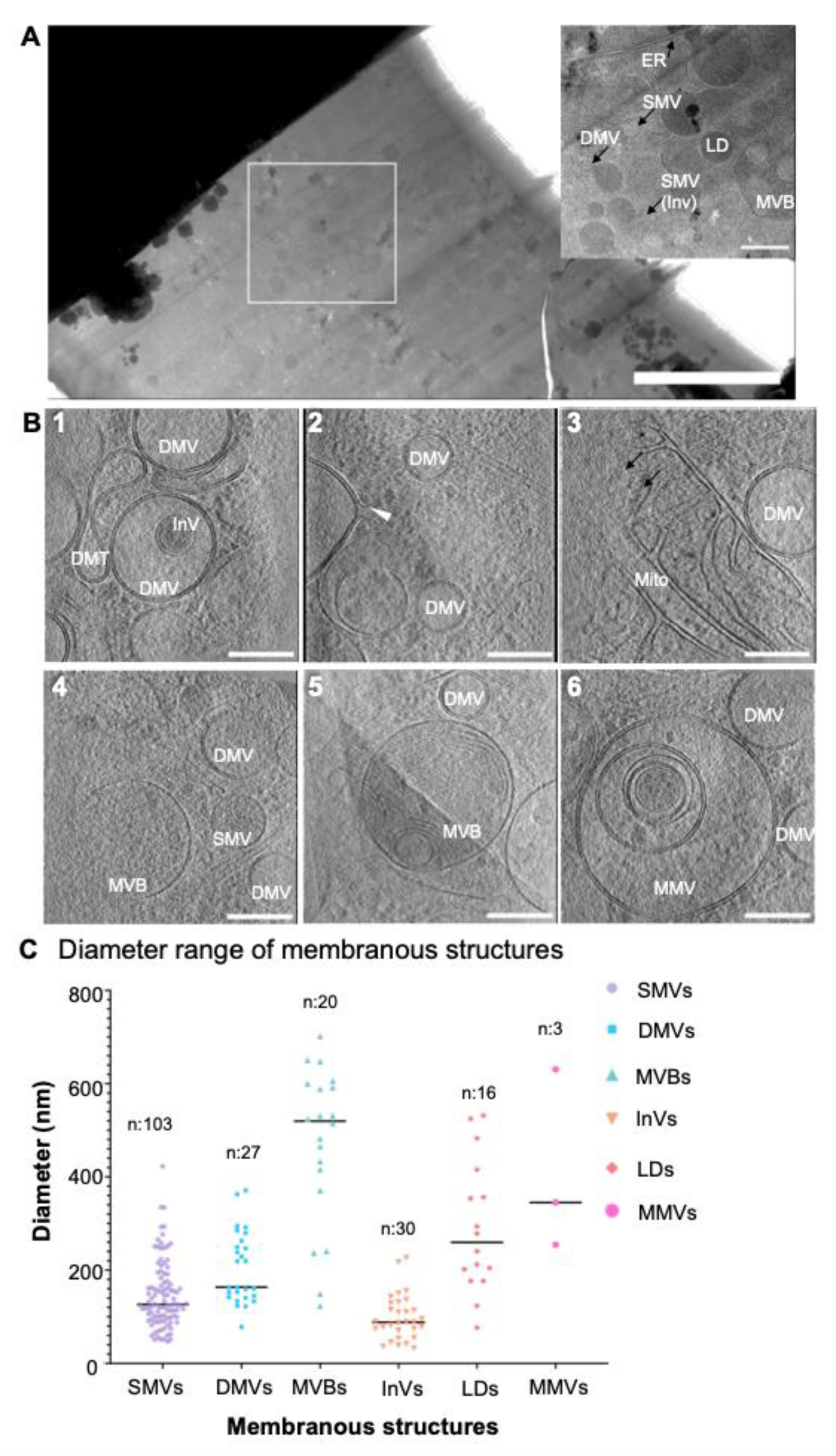
HCV membranous structures imaged by cryo-FIB/cryo-ET of Huh7 cells harbouring SGR-NS5A-eGFP. A) Representative search map (magnification 15,000X) of a lamella containing membranous structures around LDs. An area highlighting specific structures is shown in the inset. Scale bars: search map, 2 µm; inset, 500 nm. B) Representative sections through reconstructed tomograms showing the architecture of HCV MW. Scale bars: 200 nm. The white arrow indicates the opening of a DMV towards the cytosol. C) Graph depicting the diameters of the 199 observed membranous structures. SMV: Single membrane vesicle. DMV: Double membrane vesicle. MMV: Multi membrane vesicle. InV: Inner vesicle. LD: Lipid droplets. MVB: Multivesicular bodies. Mito: Mitochondria.

Based on the hypothesis that DMVs are the main sites of replication (Romero-Brey et al., 2012; Romero-Brey and Bartenschlager, 2015), the initial investigation centred on studying the ultrastructure of DMVs and their surrounding environment. DMVs were often surrounded by double-membrane tubules (DMTs), SMVs, and MVBs. Most of the DMVs examined had a closed configuration (i.e. no pores connecting the inside of the DMVs to the cytoplasm were observed; Figure 3B-1), with only one instance (1 out of 112 tomograms, consisted of two open DMVs) exhibiting an aperture facing towards the cytosol (Figure 3B-2).

As an initial characterisation of the different vesicles, the diameter of all membranous structures that were clearly visible (n = 199) was measured (Figure 3C). The majority of SMVs, DMVs and inner vesicles (InVs, found within DMVs and SMVs) had a diameter between 50 and 300 nm, whilst LDs displayed a wider range of diameter, ranging from 100 to 500 nm. MVBs, the most abundant type of vesicles imaged in the datasets, were much larger and mostly above 400 nm in diameter. However, their larger size rendered them only partially visible within the tomograms, making it challenging to accurately measure their diameter. For this reason, the diameter of only 20 MVBs was measured. Finally, MMVs were scarce, observed only in 11 tomograms. However, only 3 could be measured and had a diameter similar to that of MVBs.

Next, differences in the general distribution of densities within DMVs and SMVs were examined. Most DMVs (92.8%) either contained faint densities that could be a result of the inherent noise within cellular cryo-electron tomograms or appeared empty (Figure 4A-1,2). Only 7.14% DMVs contained patches of densities on the inside of the membrane that could potentially represent an assembly of the replicase components (Figure 4A-3, 4B-1). Finally, 17.8% DMV contained an InV (either a single or double membrane, Figure 4B-1,2,3).

**Figure 4:**
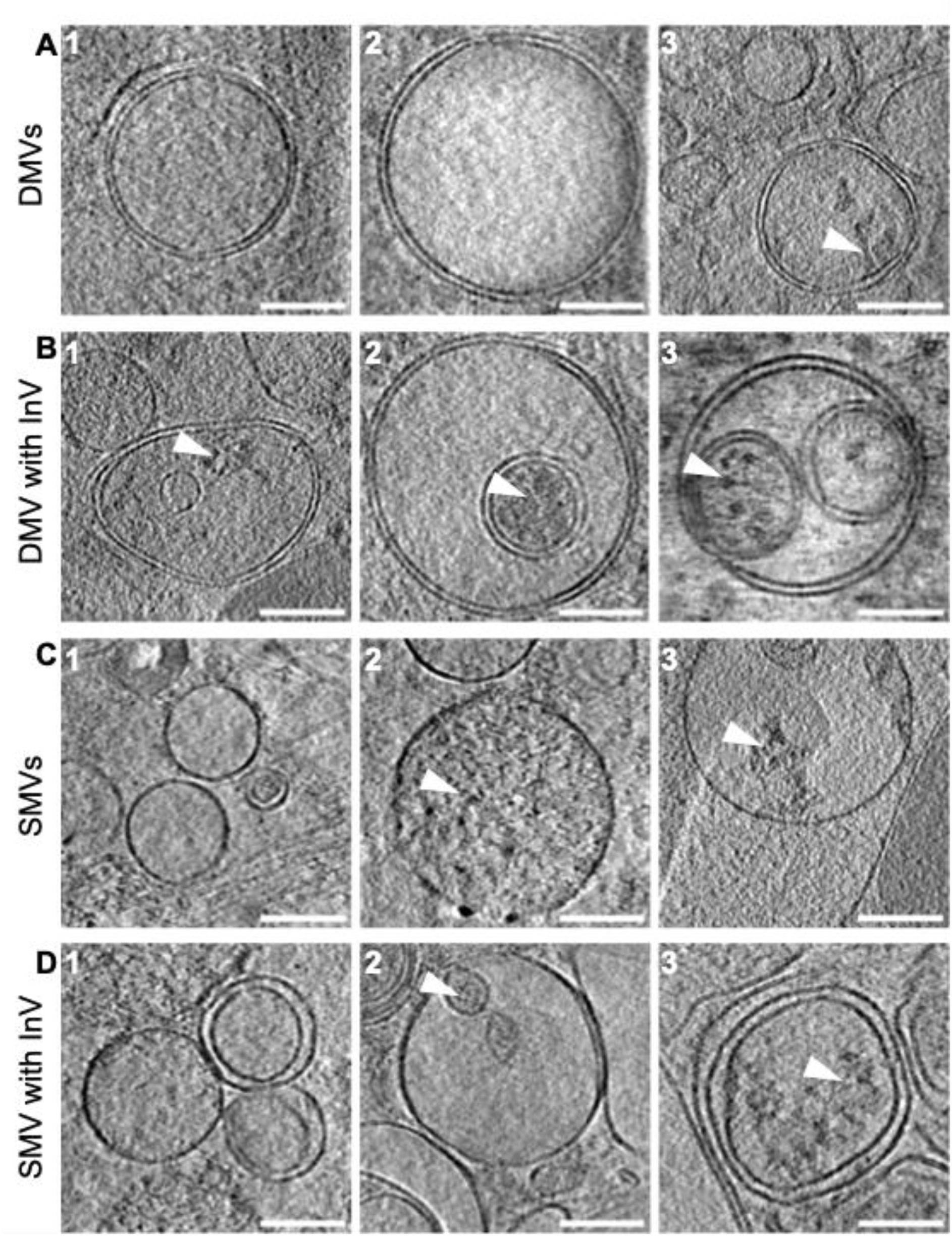
Differences in densities present within DMVs and SMVs. A) Representative tomographic slices showing the differences in the content within DMVs. B) Representative tomographic slices showing InVs within DMVs and their contents. C) Representative tomographic slices showing the differences in the content within SMVs. D) Representative tomographic slices showing InVs within the SMV and their content. Scale bars: A1,2; B2,3; C1,2 and D1,2,3: 100 nm, A3, B1and C-3: 160 nm. White arrows indicate densities present within the vesicles. SMVs: Single membrane vesicles. DMVs: Double membrane vesicles. InV-Inner vesicle

On the other hand, most SMVs (51.4%, Figure 4C-1, 4D-1) contained faint densities or appeared empty making it difficult to confirm their contents, 28.15% contained homogeneous densities throughout the vesicle (Figure 4C-2). Only 2.9% SMV contained patches of densities (Figure 4C-3), which could indicate the presence of either cellular or viral proteins. Similar to DMVs, 19.4% of SMVs contained InVs; Figures 4D-1,2,3). Out of 31 InVs measured for diameter (6 inside DMVs and 20 inside SMVs), 38% of them contained densities, 32% of them could not be confirmed but possibly have densities, 25% of them appeared empty.

Previous reports by immunogold labelling suggested that both NS3 and NS5A primarily labelled rER and SMVs with 50-70 nm diameter and to a lesser extent on DMVs (Romero-Brey et al., 2012) and thus at least some of these densities could potentially be NS3 and NS5A proteins. Overall, this analysis suggests that DMVs and SMVs are similar in terms of content, but a larger percentage of SMVs contained internal densities, which might correspond to viral and/or cellular proteins. Strikingly, the only membranous structure that contained a significant percentage of inner densities were InVs.

### NS5A-eGFP preferentially locates around MMVs

So far, a clear organisation of the replicase machinery within specific structures has not been established using the cryo-FIB milled cryo-ET datasets, due to limitations in utilising NS5A-eGFP fluorescence for guided tilt-series collection on the lamellae. Therefore, an alternative workflow was developed, incorporating CEMOVIS. This approach aimed to gather evidence regarding the types of structures close to NS5A, potentially housing the replication complex in cells harbouring SGR-NS5A-eGFP (Figure 5). An advantage of this approach is its ability to employ a heterogeneous population of SGR-NS5A-eGFP cells, thereby eliminating the need for FACS sorting. NS5A-eGFP puncta are detectable in cryo-ultrathin sections following cryo-ultramicrotomy and can thus serve as markers for guided tilt-series acquisition.

**Figure 5:**
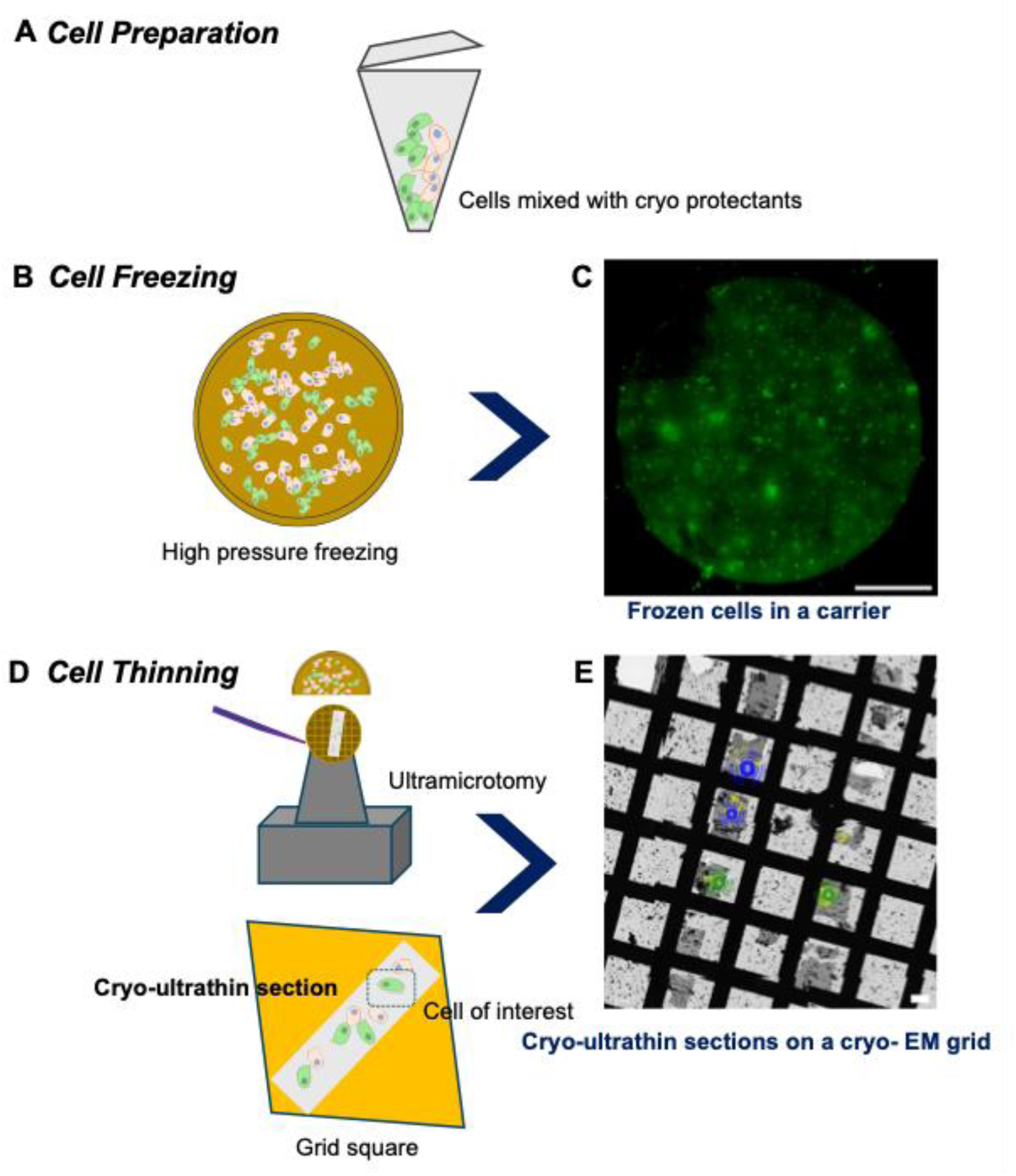
Schematic sample preparation workflow to generate cryo-ultrathin sections by CEMOVIS of Huh7 cells stably harbouring SGR-NS5A-eGFP. A) Cells were initially mixed with a cryoprotectant. B) Subsequently, cells were added to gold carriers and high-pressure frozen and imaged by cryo-fluorescence microscopy (C). Scale bar: 500 µm. Finally, cells were thinned by CEMOVIS (D) to generate ultrathin sections (E). Scale bar: 2 µm.

Using this approach, ultrathin sections of varying thicknesses (100 nm, 70 nm, and 40 nm) were evaluated, with the 40 nm sections displaying clear cellular features, whereas the thicker sections exhibited low contrast. Therefore, 12 tilt-series were collected from a single 40 nm section, guided by NS5A-eGFP foci and subsequently correlated (Figure 6A).

**Figure 6:**
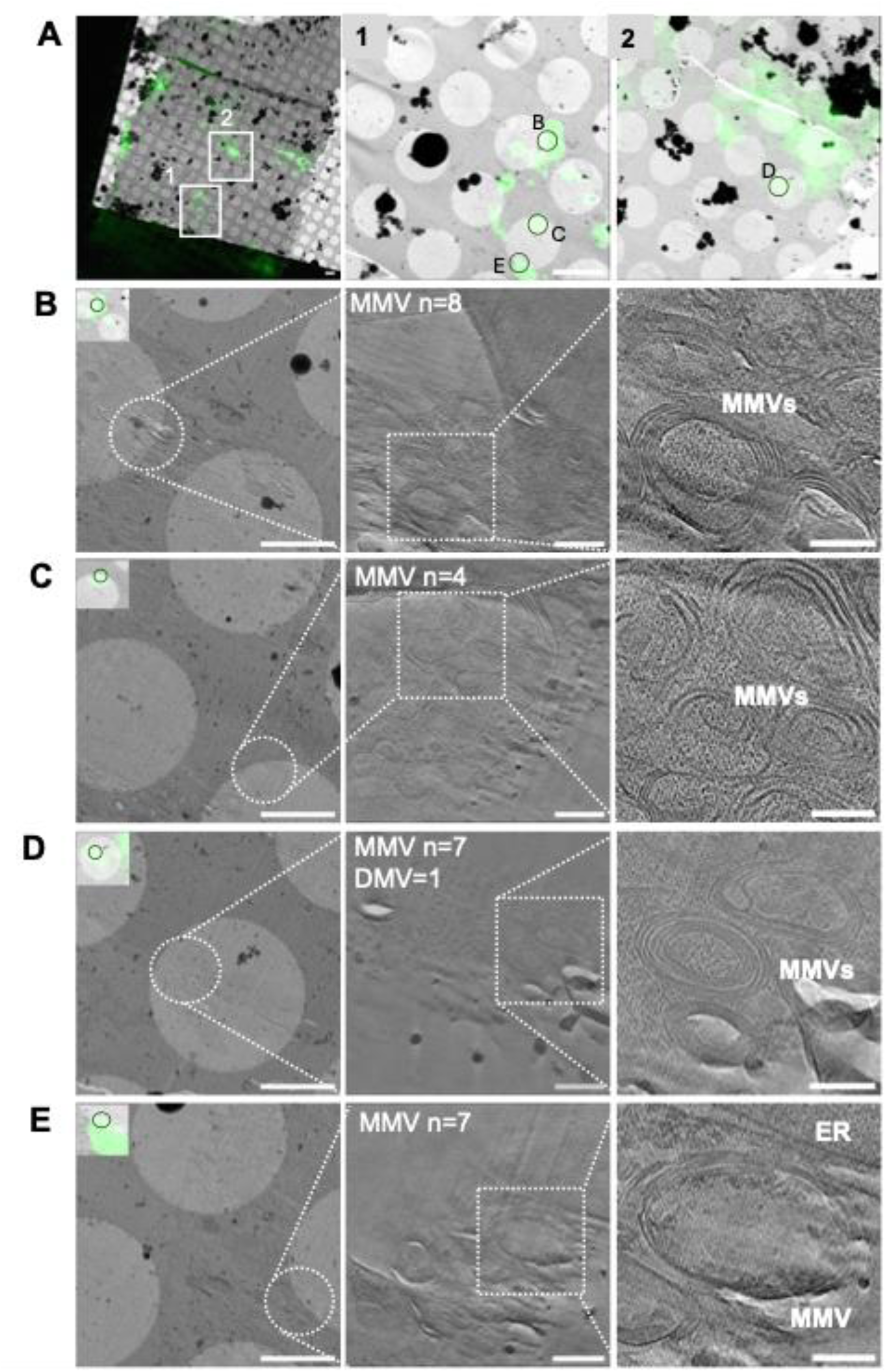
NS5A-eGFP-guided acquisition of HCV MW observed on Huh7 Cells stably expressing HCV SGR-NS5A-eGFP by CEMOVIS and cryo-ET. A) Left, low magnification search map (125 X) of a 40 nm cryo-ultrathin ribbon with regions of NS5A-eGFP signal shown in white boxes. Centre and right, the area of NS5A-eGFP foci used for tilt-series collection is shown with green circles and labelled with designated letters that correspond to the search maps and tomograms below. Scale bar: 2 µm. B-E) Left, cryo-EM search map (580 X) with highlighted position by a white circle where the tilt series was acquired. Centre and right, tomographic sections at low (centre) and high magnification (right) of membranous structures observed in the tomograms acquired at the specified NS5A-eGFP signals. Scale bars left, 100 nm; centre, 200 nm; right, 100 nm. MMVs: Multi membrane vesicles. DMV: Double membrane vesicle.

Reconstructed tomograms from 12 NS5A-eGFP tilt-series contained 83.3% MMVs, 33.3% ER, 25% DMVs, 25% mitochondria, 16.6% LD and 8.3% MVB. 12 tomograms primarily revealed 39 MMVs with notable densities inside (4 of which are illustrated in Figure 6B-E) and only 3 DMV identified. Similar densities were observed within and around DMVs as noted in the cryo-FIB-milled dataset. Interestingly, the proportion of DMVs and MMVs was reversed between tomograms obtained using the CEMOVIS and cryo-FIB workflows. In the cryo-FIB datasets, which were not guided by NS5A-eGFP presence, DMVs were more prevalent, comprising 13% of the vesicles (27 DMVs out of 199 structures measured). In contrast, MMVs dominated in the CEMOVIS datasets, comprising 80% of the vesicles (39 MMVs out of 49 structures identified), which were exclusively collected in regions exhibiting NS5A-eGFP signals. Strikingly, one of the MMVs displayed a heterogeneous arrangement of proteins, potentially indicative of a replication complex assembly (Figure 6E). Notably, this represents the most organised assembly of cryo-densities observed across all datasets, underscoring the value of fluorescence-guided tilt-series collection on thin cryo-specimens.

## Discussion

The use of direct-acting antivirals (DAAs) has dramatically improved the life of HCV-infected patients, resulting in a reduction in the global burden of HCV from 170 million individuals 10 years ago to the current estimate of 50 million (WHO, 2024). The targets for DAAs (NS3 protease, NS5A and NS5B RNA-dependent RNA polymerase) are all directly involved in virus genome replication. It is thus important to understand the molecular details of this process as this will shed light on the mode of action of DAAs and may contribute to an understanding of DAA resistance which is becoming increasingly common. In this regard one important avenue of research has focused on understanding how HCV modifies the host membranes to establish its replication complex. From 2002 to 2019 a number of research groups explored the membranous structures within the MW in relation to HCV infection or the presence of SGR, using a mix of light and electron microscopy techniques (Ferraris et al., 2010; Romero-Brey et al., 2012; Paul et al., 2014; Berger et al., 2014; Mohl et al., 2016; Pérez-Berná et al., 2016; Lee et al., 2019). However, these studies relied on chemical fixatives and resin-embedded sections, limiting direct visualisation of key replication components such as dsRNA and protein arrangements within membranous structures. Furthermore, even though DMVs have been identified as the most probable structures to harbour the replication complex (Romero-Brey et al., 2012), the location of the replication complex is still under debate based on direct visualisation of assembly of viral and cellular proteins. Here, we aimed to develop a cryo-ET workflow to visualise HCV MW in close-to-native conditions in the absence of chemical fixation, and to explore the localisation of the HCV replication complex within it. A detailed workflow was developed, spanning from sample preparation to data analysis, incorporating the latest cryo-ET techniques, and two cryogenic sample preparation techniques were established for an *in-situ* investigation of the membranous structures in cells containing an SGR (HCV genotype 2a): cryo-FIB milling (Lam and Villa, 2021) and CEMOVIS (Chlanda and Sachse, 2014).

HCV-induced host membrane rearrangements are driven by viral proteins. In SGR harbouring cells host membrane modifications similar to those in HCV-infected cells were observed (Romero-Brey et al., 2012). To track viral proteins, we utilised an SGR-NS5A-eGFP replicon. This construct is the only HCV system with a fluorescently tagged non-structural protein. However, one limitation of this approach is that NS5A is involved in both replication and assembly (Eyre et al., 2014), suggesting that some observed NS5A-eGFP signals may not be exclusively associated with replication sites.

Datasets acquired using cryo-FIB milling followed by cryo-ET (Wagner et al., 2020) without fluorescence guidance enabled the exploration of the MW induced by the SGR-NS5A-eGFP in Huh7 cells. Analysis of these datasets revealed numerous DMVs and SMVs. While most DMVs appeared empty or contained densities at background levels, SMVs frequently exhibited either patchy densities or were entirely filled. Additionally, both DMVs and SMVs occasionally contained inner vesicles (InVs) with densities present within. These InVs have been previously described as self-invaginations of SMVs (Romero-Brey et al., 2012) or vesicles in cluster (ViCs) (Ferraris et al., 2013). However, they may also represent exosomes containing dsRNA or non-structural proteins that fuse with larger vesicles, potentially serving as replication sites (Ramakrishnaiah et al., 2013; Bukong et al., 2014; Yin et al., 2022). Given that positive-sense RNA virus replication sites are expected to contain viral replicase and RNA (Wolff et al., 2020), our findings suggest that SMVs and InVs may serve as primary sites for HCV SGR replication. This is further supported by EM studies of chronically infected HCV patient cells, which identified SMVs and not DMV around LDs and the ER (Blanchard and Roingeard, 2018).

In any case, the most frequently observed vesicles in the cryo-FIB/cryo-ET dataset were the MVBs, consistent with EM analysis of replicon cells transfected with NS5A-mCherry (Grünvogel et al., 2018), suggesting their involvement in the process of replication. The aggregation of MVBs, found primarily near SMVs, DMVs, and LDs, with diameter sizes ranging from 400–800 nm (though most exceeded 1000 nm) were previously described as single-membrane compartments containing multiple circular units with dense cores (Ferraris et al., 2010). The presence of densities within SMVs and DMVs within MVB also aligns with the previous observations of NS5A-mCherry localised inside MVBs (Grünvogel et al., 2018) and with the dense lumen within concentric units (Ferraris et al., 2010). The smaller SMVs (30–150 nm) observed within MVBs resemble exosomes in size. Previous studies have demonstrated that exosomes derived from HCV-infected cells contain viral RNA, proteins, and complete virions (Yin et al., 2022). Notably, these exosomes facilitate the transfer of HCV RNA to uninfected cells, even in the presence of neutralising antibodies (Ramakrishnaiah et al., 2013). This suggests that MVBs play a role in HCV RNA dissemination. On rare instances, tomograms captured SMVs invaginating into MVBs (not shown), supporting the hypothesis that MVBs may transport vesicles containing replicase machinery or dsRNA to neighbouring cells (Grünvogel et al., 2018). Perhaps, the accumulation of MVBs in SGR harbouring cells is due to the lack of viral assembly in this system, thus depriving nascent genomic RNA of a ‘final destination’? Nonetheless, this observation suggests a potential role for MVBs in HCV replication Previous immunofluorescence studies indicated that HCV replication vesicle membranes originate from the ER but also colocalise with markers of early and late endosomes, coat protein complex (COP) vesicles, mitochondria, and LDs (Romero-Brey et al., 2012), as well as lysosomes (Matsui et al., 2021). The measured SMV diameter (50–300 nm) aligns with that of endosomes (100–500 nm) and lysosomes (200–300 nm). If these SMVs are derived from early endosomes, their internal densities may represent NS3 and NS5A, while surface-exposed densities may correspond to GFP-Rab21 (Romero-Brey et al., 2012). Alternatively, if they originate from lysosomes, internal densities may include NS5A and LAMP-2A (Matsui et al., 2021).

Notably, in CEMOVIS datasets guided by NS5A-eGFP fluorescence, MMV accumulation was observed. One tomogram revealed densities arranged in a regular pattern, potentially representing replicase machinery assembly on the inner leaflet of an MMV. Overall, this study contributes to the development of cryo *in-situ* workflows for investigating HCV-induced membranous structures. Our findings show that DMVs appear mostly empty, while SMVs and InVs contain significant densities. Additionally, MVBs emerge as the most abundant vesicles in MW imaging at random, reinforcing their potential role in HCV replication and RNA transfer. The most abundant vesicles among NS5A-eGFP fluorescence-guided cryo-ET were MMVs, suggesting that are involved in replication in cells harbouring SGR-NS5A-eGFP.

In conclusion, these findings provide new insights into the spatial organisation of NS or cellular proteins within membranous structures in replicon-transfected cells, revealing structural details previously inaccessible through conventional electron microscopy. As cryo-ET continues to evolve with advancing computational techniques, it will further refine our understanding of NS protein interactions with cellular proteins, and HCV RNA genome organisation within these membranous structures.

## Acknowledgements

UMS was supported by a Wellcome PhD studentship (222370/Z/21/Z). HCV studies in the MH laboratory were supported by a Wellcome Investigator Award (096670/Z/11/Z) and an MRC project grant (MR/S001026/1). JF was supported by grant PID2023-149259NB-I00, funded by MICIU/AEI/10.13039/501100011033 and by “ERDF A way of making Europe”. The funders had no role in study design, data collection and analysis, decision to publish, or preparation of the manuscript.

We acknowledge the technical contributions of the University of Leeds Bioimaging Facility, especially Dr Ruth Hughes and Dr Sally Boxall, and the Astbury Biostructure Laboratory Electron Microscopy facility, especially Mr Martin Fuller, Dr Rebecca Thompson; and the Electron BioImaging Centre (eBIC) at Diamond Light Source, especially Dr James Gilchrist.

## Author Contributions

UMS, JF and MH planned the study. UMS, TJO’S, and YH performed the experiments. UMS and JF analysed the data. UMS, JF and MH wrote the main manuscript text. All authors reviewed the manuscript.

## Data availability

The datasets generated during and/or analysed during the current study are available from the corresponding authors on reasonable request.

